# Griottes: a generalist tool for network generation from segmented tissue images

**DOI:** 10.1101/2022.01.14.476345

**Authors:** Gustave Ronteix, Valentin Bonnet, Sebastien Sart, Jeremie Sobel, Elric Esposito, Charles N. Baroud

## Abstract

Microscopy techniques and image segmentation algorithms have improved dramatically this decade, leading to an ever increasing amount of biological images and a greater reliance on imaging to investigate biological questions. This has created a need for methods to extract the relevant information on the behaviors of cells and their interactions, while reducing the amount of computing power required to organize this information. This task can be performed by using a network representation in which the cells and their properties are encoded in the nodes, while the neighborhood interactions are encoded by the links. Here we introduce `Griottes`, an open-source tool to build the “network twin” of 2D and 3D tissues from segmented microscopy images. We show how the library can provide a wide range of biologically relevant metrics on individual cells and their neighborhoods, with the objective of providing multi-scale biological insights. The library’s capacities are demonstrated on different image and data types. This library is provided as an open-source tool that can be integrated into common image analysis workflows to increase their capacities.

## I. INTRODUCTION

The effect of the cellular heterogeneity on the spatial organization and function of biological tissues is emerging as a major paradigm to understand complex collective behaviors. This is the case for example in the study of how spatial structure leads to the emergence of germ layers during embryonic development [1] or in describing the tumor heterogeneity in cancer [2–5]. Indeed modern *in vitro* models such as in organ-on-a-chip devices [6] or organoids [7] capture complex biological phenomena by integrating several cell types that follow a precise spatial patterning. These models reinforce the view that interactions between cells are fundamental in determining many properties of the tissues. In particular they show how the mechanical and biochemical interactions between neighboring cells determine many global biological properties across many spatial scales [4].

In this context fluorescence microscopy has emerged as an invaluable tool to generate large and high-resolution images of two-dimensional (2D) or three-dimensional (3D) tissues. These images have a typical resolution below one micron, with several fluorescence channels being acquired in parallel, often on several planes. As a result it is now common to generate thousands or more multi-channel images that represent a trove of data on the tissues being imaged (see e.g. Ref. [8]). This trend is amplifying, with the increase in performance of microscopes, cameras, and fluorophores, such as e.g. the new CODEX technology [2, 9] that makes it possible to simultaneously image close to 60 markers in tissues. These advances in image acquisition have led to a need for analysis methods to extract biologically relevant information from these images, namely at the scale of the individual cells. Such methods need to preserve information about the cell types, their positions, and their biological function, while reducing the size and complexity that make the original images difficult to treat.

Geometric networks constitute a natural representation of spatially encoded information that can be used to describe the structure and interactions in biological tissues. The network representation has already been used in a range of settings, ranging from city structure and railways [10] to flows in the pulmonary airway tree [11]. In contrast with protein interaction networks that are common in biology, geometric networks preserve the spatial information of the sample while encoding the properties of the individual objects. This makes them well-suited for describing the structure and interactions between cells and to bridge the scales from the individual cell to the tissue. More fundamentally, representing tissues as networks is advantageous as it is possible to draw from a vast literature in statistical physics, mathematics and computer science to analyze the structure and quantify the relationships between the cells composing the tissue [12].

Here we describe `Griottes`, an open-source tool to construct a network representation from imaging data. The routine creates a “network twin” of the imaged sample by extracting individual cell information from segmented images and building a network representation of the tissue. This new object makes it easier to address quantitative questions on complex tissues with single-cell precision. Below we begin by describing the code and its function, followed by a a few examples on how biologically relevant information can be obtained from typical images.

## II. RESULTS

A network is a set of objects that are related to each other. Each object is described by a node, which can have several attributes, and the nodes are connected by edges or links that represent the relationship between them. The network is represented by a graph that shows these objects as nodes and their relationships as links. In `Griottes` we choose to represent each cell by a node that is placed on its geometric center in 2D or 3D. The links in the network represent neighborhood, either through contact or through distance, as described below. Different attributes (cell type, cell size etc.) can be assigned to the nodes, while the links can contain information about the distance between cells or the length of the common border, as summarized in Table. I.

**Table I.** List of attributes and their definitions as currently available in `Griottes`.

| Node Attribute | Definition |
| --- | --- |
| x, y, z | Mask geometric center in $x, y, z$ |
| area | Mask size |
| $\theta$ (, $\psi$) | Orientation of the mask major axis (in 3D) |
| eccentricity | Mask eccentricity |
| fluorescence | Mean mask fluorescence (one value for each channel) |
| label | Unique number each cell |
| Link attribute | Definition |
| Contact size | Length of the cell-cell contact |
| Distance | Distance between two cell centers |

`Griottes` takes as an input segmented tissue images and builds a the graph to represent them. The input to `Griottes` can consist of data tables, as well as labeled images of nuclei and/or cytoplasms. The inputs can be 2D or 3D, without any impact on the program and a single line of code is enough to generate the graph representation of the tissue.

### A. Program architecture

The `Griottes` analysis pipeline contains two main steps: individual cell attribute measurements step and then the network construction step. These processes and the program inputs and outputs, described in figure 1, are elaborated immediately below.

**Figure 1.**
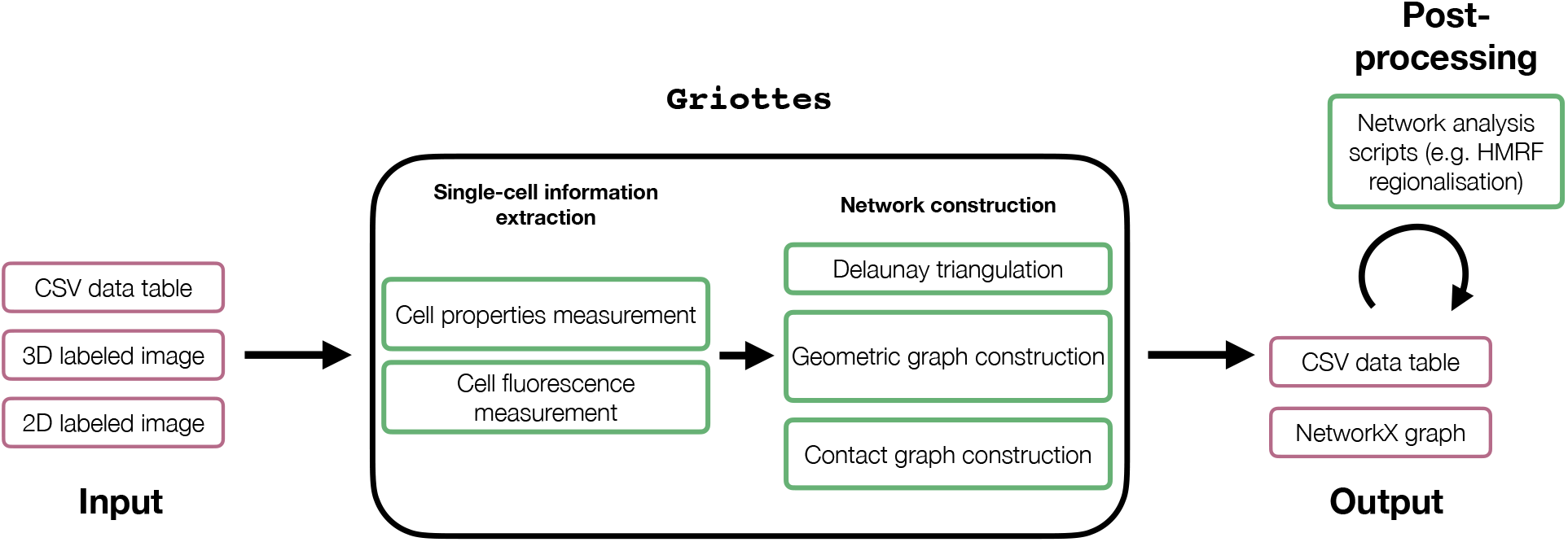
Schematic representation of the `Griottes` workflow: the program takes multiple data formats as input to generate a graph in a single command. The red boxes contain objects, the green boxes processes.

#### a. Input

`Griottes` takes as input the output of standard cell segmentation codes. This can be in either as a labeled image (2D or 3D) or as a data table. The labeled image has at least one channel composed of a zero-padded matrix in which the area occupied by each cell is filled with an integer label that is unique to this cell. A data table containing single cell information can also be used as an input (for supplementary details on the input formatting we refer the reader to the program documentation).

#### b. Individual attribute measurements

If the input is an image containing segmented cells, `Griottes` can extract their essential geometric properties such as their size, shape and position in the image (see table I). In case the input image also contains supplementary fluorescence channels, then the individual fluorescence intensities of the different cells can be extracted too. The data are returned to the user, giving access to the statistics of every single cell composing the tissue.

#### c. Network construction

From the individual cell properties, `Griottes` builds a spatial graph where each node can be populated with cell attributes defined by the user. The construction methods are elaborated below.

#### d. Output

The returned object is a graph where the previously measured properties are entered as node attributes, each node in the network mapping to a given cell in the tissue. The object is currently returned as a NetworkX graph [13].

#### e. Post-processing

After the network image of the tissue has been generated with `Griottes`, it can be further analyzed downstream using tailor-made programs depending on the biological question to be addressed. Example applications include spatial clustering algorithms, such as Hidden Markov Random Fields (HMRF), or graph neural networks (GNNs) to investigate the neighborhood influence on biological properties such as cell division [14]. Once the network image has been generated, it is possible to harness the diverse graph analysis methods to extract relevant information from the experimental data (Fig. 1).

### B. Analysis pipeline

#### 1. Extracting single-cell information from the image

The input image has to contain at least a labeled image that contains the information on the segmented cells. This allows the conversion of pixel-based information on the different cell types to the node-based representation of the tissue. In the following paragraphs we summarize the different features that can be extracted using `Griottes`.

##### a. Geometric property measurements

The geometric properties can be of particular interest as it is known that cell morphology can be a significant feature of tissues and their characterization can shed light on morphogenesis or mechanical constraints within the tissue [15]. To be able to readily include the morphological information as a feature of the cells in the network, we added an option to measure the geometric properties of the cell masks in `Griottes`. This includes extracting the mask volume, eccentricity and orientation in space.

##### b. Fluorescence measurements

If the input image contains supplementary channels containing the fluorescence information, `Griottes` can measure the mean fluorescence levels inside each cell mask and attribute this number to the cell. This allows for downstream cell classification (is the cell of type *A* or *B*?), or phenotype information (what is this cell’s production of molecule *C*?). The method is particularly reliable when the cell masks are built from the membranes and the full volume of the cell is recovered. If only the nucleus is stained, then only the information from the pixels inside the nucleus mask is taken into account.

#### 2. Building the network

Once the information on the cell properties is extracted from the image, `Griottes` constructs the connectivity graph binding neighboring cells together (Fig. 1). Several options are implemented by the program for determining the connectivity between two cells.

The first method uses geometric graph construction. A geometric graph is generated by connecting cells separated by less than a given distance [10]. This connection method can be particularly relevant when cell-cell communication is mediated by chemical cues [16], in which case the distance between cells determines the time necessary for diffusion to take place. However, this construction rule is less pertinent if cell-cell interactions are contact mediated.

The second method is a Delaunay-triangulation based connection condition (Fig. 2b). This is a robust state-of-the-art heuristic widely used to build biologically relevant networks in many different contexts [16–18]. The Delaunay triangulation provides a good estimate of the cellcell connectivity in the cases when the membranes are not available or cannot be identified in the image data. However care must be taken when analyzing such results, since this method is known to not perfectly replicate tissue morphology in 2D [19]. In order to avoid incoherent links generated by the Delaunay procedure, the connections between cells beyond a cutoff distance are removed from the network.

**Figure 2.**
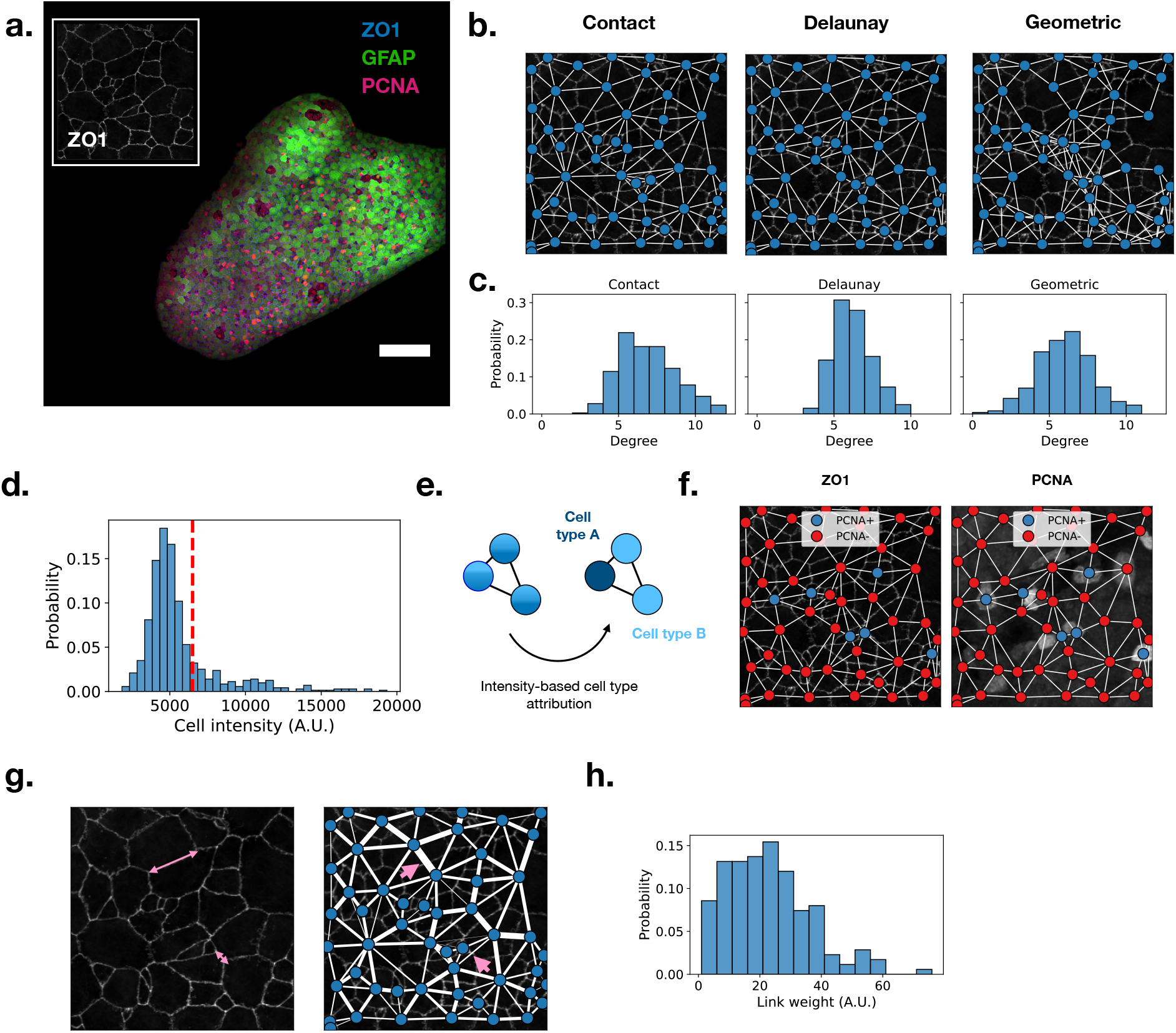
**a.** Confocal dorsal view of the zebrafish adult pallium. GFAP in green, PCNA in magenta and ZO1 in white or blue. ZO1 is highlighting the apical domain of the cells allowing the identification of their apical area. Inset: spotlight on a limited tissue area, the ZO1 membrane staining allows for the exact localization of cell membranes. Scale bar is 100*μ*m. **b.** `Griottes` incorporates different network construction methods. The contact-based method connects cells sharing a common membrane, the Delaunay and Geometric graphs are commonly used graph-generation methods. **c.** The graphs generated from a same set of nodes with different construction rules have different properties. For instance, the degree distribution of a Geometric graph is broader than that of the Contact and Delaunay graph. **d.** Mean PCNA signal within the cells in the example tissue. Cells with an average intensity above 6500 (red line) are considered PCNA+, the other cells are PCNA-. **e.** Thresholding intensity signals converts a network populated with continuous fluorescence signals to a network populated with categorical cell types. **f.** Representation of the example network (panel **b.**) where node colors represent cell type. Left: the network is projected on the ZO1 signal. Right: the network is projected on the PCNA signal. This method reliably incorporates cell type information into the network representation of the tissue. **g.** Left: connected cells can have widely varying contact surfaces. Right: this information can be encoded into the network by weighting the links between cells. Two differing cell-cell interfaces (pink lines) have different link weights in the network representation of the tissue (pink arrows). **h.** The connection between cells can be quantified at scale: the histogram of link weights in the tissue.

In contrast with the above cell-center based methods, if the segmentation recovers masks of the complete cells, it is also possible to construct a graph that accounts for direct contact between the cells. This method relies on the interfaces between the different cells (Fig. 2b), linking cells that share a common membrane. With this method it is possible to provide values to the edges as a function of the shared membrane surface, as described in Table I. This feature can be transposed in the graph object output by weighting the edges that link two nodes (Fig. 2g).

## III. APPLICATIONS

The method and interest in using network description of tissues is clearer when the library is applied to specific cases. Below we provide some illustrations of the implementation of `Griottes` on three types of inputs: 2D and 3D images as well as a data table. These examples also correspond to different biological samples: the adult zebrafish pallium and mesenchymal stem cell spheroids. Each example showcases a different use case for the program.

### A. Zebrafish telencephalon 2D image

Deciphering the generation of neural stem cells (NSCs) in the adult zebra fish brain requires a deeper understanding of the connections between individual cells in the tissue. High-quality *in vivo* images have recently been obtained by Dray et al.[20], as shown in Fig. 2a. This image consists of a projected 3D stack of whole-mount immunostained pallium from 3 months post-fertilization zebrafish. Visualization of GFAP (glial fibrillary acidic) expression allows the identification of the neural stem cells, while PCNA (proliferating cell nuclear antigen) highlights dividing cells and ZO1 (Zonula Occludens 1) shows the apical domain of the cells. These cells have been shown to coordinate their behavior in part via local cell-cell interactions (see Ref.[20] for details).

The image shown in Fig. 2a is first segmented using Cellpose [21], which yields a multi-channel 2D image containing the masks resulting from the cell segmentation and the fluorescence image in the GFAP and PCNA channels. This image is then analyzed using `Griottes` to generate a network representation of the tissue, as shown in Fig. 2b. Different network construction methods generate different graphs. This feature is illustrated in Fig. 2b, where a graph is constructed for a sub-section of the tissue using the contact, the Delaunay and the geometric construction rules. For the geometric and Delaunay graphs, the maximum cell-cell distance authorized is 15 *μ*m, to avoid generating spurious connections between cells. While the Delaunay and contact graphs give broadly similar structure, the geometric graph shows clear differences in neighborhood, due to the heterogeneous sizes of the cells and their arrangements. The differences in graph structure can be measured quantitatively, e.g. by observing the degree distribution (number of links per cell) as a function of the network construction method. The distribution of degrees in the geometric graph is much broader than in the contact of Delaunay graphs (Fig. 2c).

From the input image, `Griottes` can also extract the fluorescence in the PCNA channel of the individual cells in the tissue. The distribution of fluorescence intensities is shown in figure 2d. A threshold value is then selected manually in order to classify cells with a higher intensity as cell type PCNA+ and those below as cell type PCNA- (Fig. 2e). This binary classification now allows us to set the cell type as an attribute for each node and construct the network representation of the tissue (Fig. 2f). We verify that the brightest cells in the PCNA channel are indeed classified as PCNA+ (Fig. 2f).

In images where the whole cells are segmented, as is the case here, it is possible to include the individual cellcell contact surfaces in the network representation of the tissue. Indeed, cells can have different shapes leading to a high heterogeneity of cell contact surfaces within the tissue (Fig. 2g). Cell-cell contact can be pertinent to understand tissue behavior, since both chemical and mechanical signals are transmitted through membrane contacts. Extracting properties of these contact surfaces can shed light on key biological processes such as notch signaling [22]. `Griottes` allows easy retrieval of the contact length between all pairs of neighboring cells. The size of the contacts can be represented visually for each link (Fig. 2g) and the quantitative distribution of these values can be obtained (Fig. 2h) for downstream analysis.

### B. Analysis of 3D spheroid images

The network representation can also be applied to 3D cultures, in the form of spheroids or organoids. These model systems recapitulate many aspects of the *in vivo* tissue structure, as shown e.g. by comparing cryosections of organs with organoids [7, 23]. In these tissue models the structural organization in 3D was shown to couple back on the functional behavior of the cells within the organoid. This was quantitatively shown in the case of mesenchymal stromal cells (MSCs), which were found to organize in a hierarchical manner, depending on their level of commitment, and to express different levels of chemokines depending on their position in the organoid [24, 25]. Previous analysis however was limited to coarse-grained quantities on averaged 2D images [26, 27]. This can now be overcome by imaging the organoids using a light-sheet microscope, which provides very good resolution over the complete depth of the organoid. Some representative slices of the 3D reconstruction are shown in Fig. 3a.

**Figure 3.**
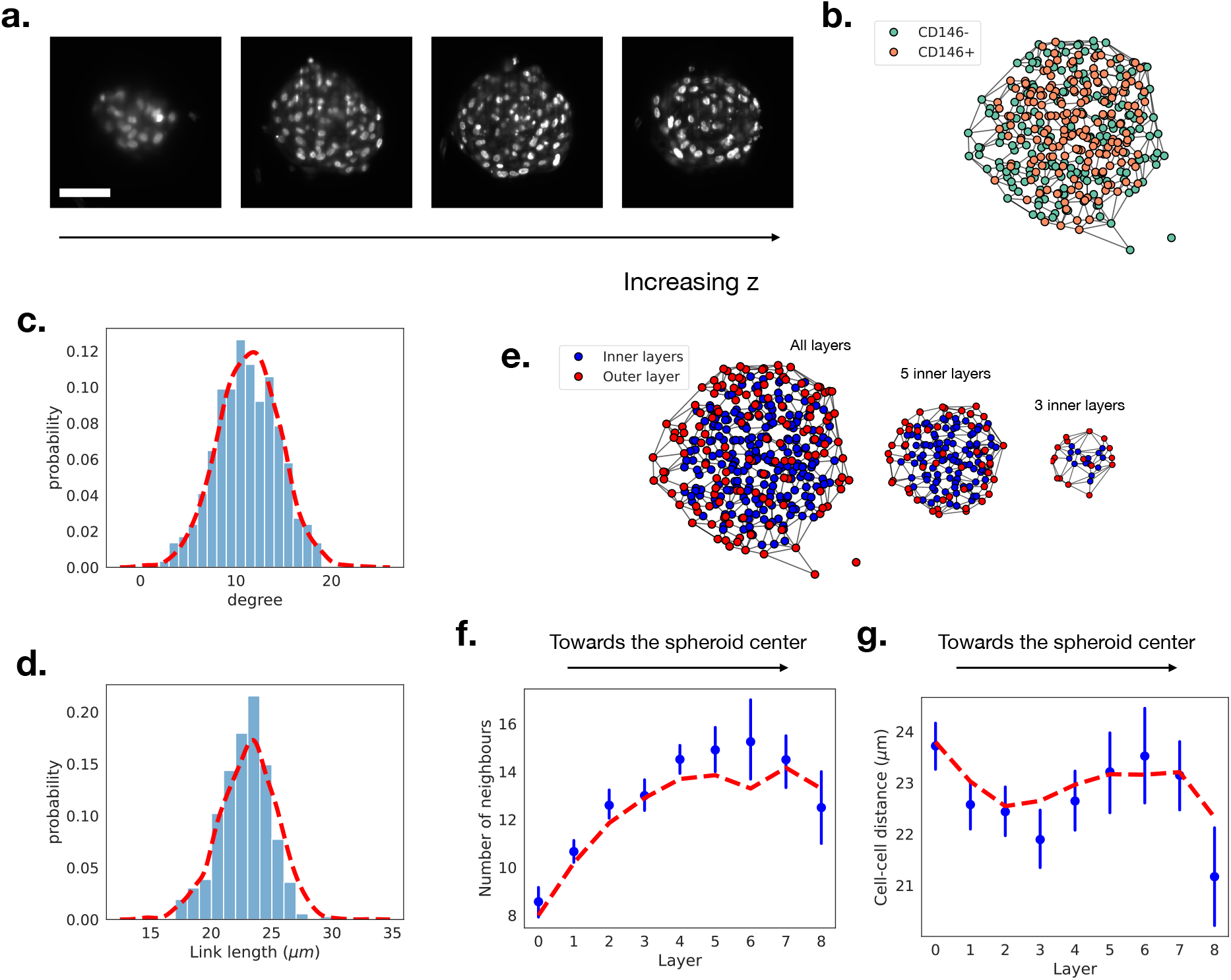
**a.** Sections of a MSC spheroid imaged with a light-sheet microscope. The technique allows in-depth imaging of tissue structures. Scale bar is 50*μ*m. **b.** 3D network representation of a MSC spheroid. Different cell types are identified based upon CD146 fluorescence measurements. **c.** Comparison between the degree distribution of an example spheroid (panel **b.**, bars) and the batch distribution (N = 5, red line). **d.** Comparison between the link-length distribution of an example spheroid (bars) and the batch distribution (red line). **e.** The network representation makes it possible to identify cells on the outer layer of the spheroid (red) from the inner cells (blue). We can “peel off” the outer layers successively, revealing the inner structure and composition of the spheroid. **f.** Cell degree as a function of the layer number, the average degree is larger for the layers near the center of the spheroid. Blue dots show one example spheroid and red dashed line represents average over the experimental batch. **g.** Distance between cell centers (in *μ*m) as a function of the layer number. Blue dots show one example spheroid and red dashed line represents average over the experimental batch.

The MSC spheroids, with CD146 and DNA staining (see the methods section for the full protocol), were imaged in 3D thereby providing access to the cell nuclei and to the distribution of CD146-expressing cells. The images were then segmented using Cellpose [21] and fed into `Griottes` in order to generate the 3D network representation of the organoid for further analysis. Since only the nuclei were available in these images, the Delaunay construction method was used to estimate the 3D graph, as shown in Fig. 3b. For illustration purposes, the cells here are classified depending on the CD146 expression with the same methodology as in section III A.

This network representation allows us to retrieve the distribution of number of neighbors (degree) for each cell within the tissue. By repeating the measurement on *N* = 5 spheroids, the distribution of degrees is found to be well-described by a normal distribution, centered at 〈*k*〉 = 11.4 ± 3.2 neighbors per cell (Fig. 3c). Other properties, such as the distribution of distances between every two neighbors within the samples, can readily be obtained as well (Fig. 3d). In the MSC spheroids, cells are on average separated by 〈*l*〉 = 23 ± 2.4 *μ*m (Fig. 3d).

It is also possible to assign a layer number to each of the cells in the network and to analyze the cells based on their distance from the edge or center of the organoid. This allows us to virtually peel off the cells composing the outer layers of the spheroid or to treat each layer individually (Fig. 3e). Analysis of the average degree of cells composing each layer, see figure 3f, shows an increase from an average of 〈*k*_0_〉 = 8.0 ± 2.4 neighbors per cell in the outer layer (layer 0) to 〈*k*_6_〉 = 13.3 ± 2.9 neighbors in the inner layers (layer 6 in this case). In contrast to the increase in the number of neighbors as a function of the layer, the cell-cell distance remains stable across the spheroid depth. Indeed, the mean distance, defined by length from one nuclei barycenter to another, varies between 〈*l*_3_〉 = 22.6 ± 2.3 *μ*m and 〈*l*_0_〉 = 23.8 ± 2.6 *μ*m (Fig. 3g). In the future, the layer analysis framework could be combined with categorical information on the cells, such as their phenotype, to assess the formation of the spheroids and organoids in greater detail.

### C. Building a network from a data table

Image processing software can often yield cell information in a table format containing the cell positions as well as other attributes such as the cell type. The data table can contain the positions of the cells in the *x–y* plane, as well as further information about the cells, such as their cell type. Below we show how such data can be integrated and analyzed using `Griottes`. In the current example a region of interest of the zebrafish telencephalon was segmented and analyzed [20], attributing a cell type to each of the visible cells in the sample. Understanding cellular organization and cell-cell relationships of this tissue will be critical to better understand how this stem cell population coordinates its behavior to sustain large-scale and long term homeostasis.

The cell positions can serve to place them on a two-dimensional scatter plot where the different cell phenotypes are indicated by different colors (Fig. 4a). `Griottes` is then used to build the network representation of the tissue, here using the Delaunay triangulation to connect cells to each other (Fig. 4b).

**Figure 4.**
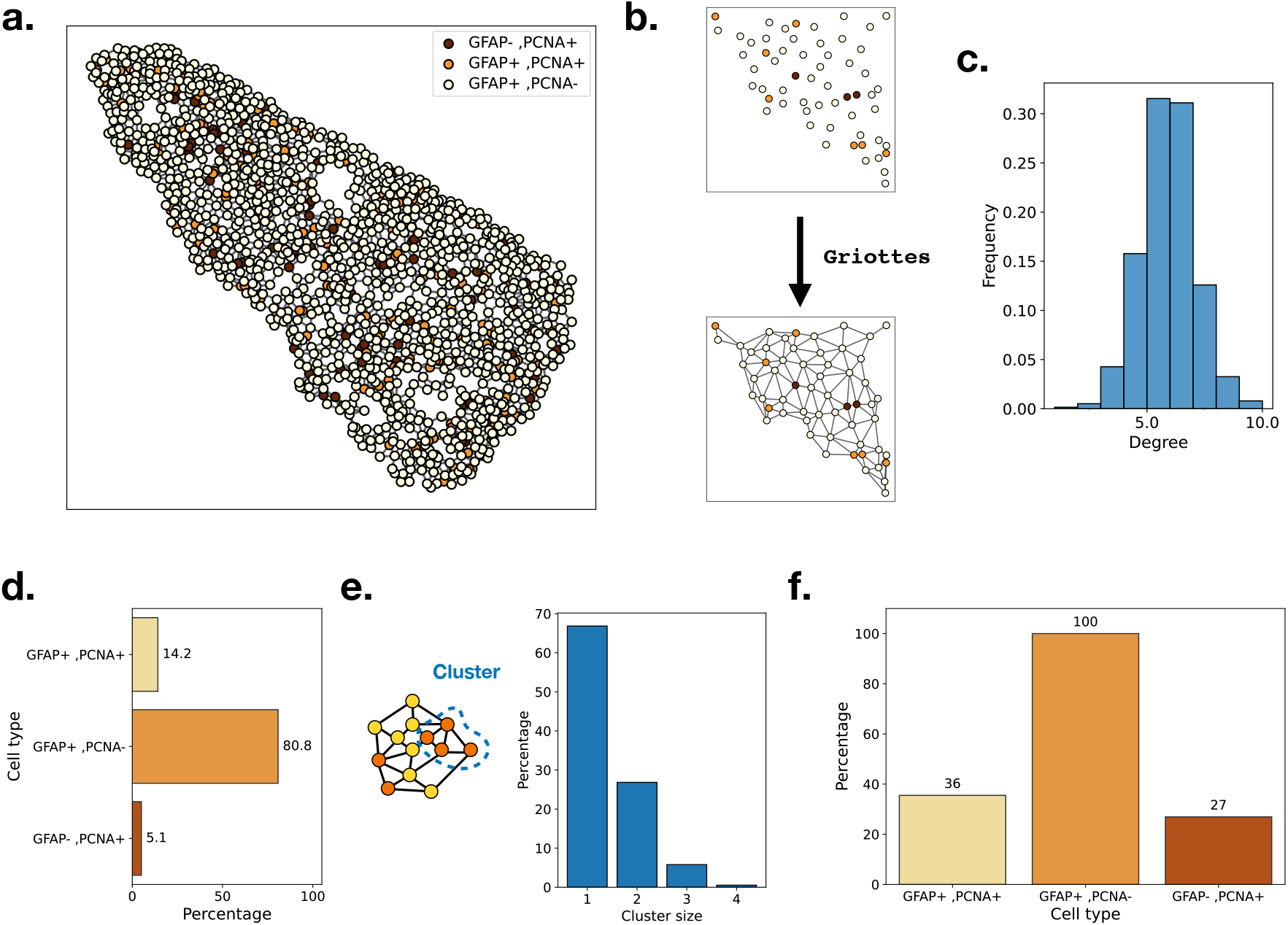
**a.** Reconstruction of the zebrafish telencephalon from a data table. Colors represent the different cell types entered in the table by the user. **b.** Network construction with `Griottes` from a point-cloud using the Delaunay construction rule. **c.** Degree distribution of cells composing the zebrafish telencephalon after the network construction using the Delaunay rule. **d.** Composition of the zebrafish telencephalon: a vast majority of the cells are GFAP+/PCNA-. **e.** From the network representation of the tissue we can extract clusters of any given cell type (left). Distribution of PCNA+ cluster sizes (right). **f.** Percentage of cells that belong to a cluster of their cell type of size larger than 1. All GFAP+/PCNA-cells are connected and belong to the same cluster. Conversely, a majority of GFAP+/PCNA- and GFAP+/PCNA+ cells aren’t connected to any other cell of the same type.

The telencephalon is structured in a quasi-2D structure, the cells spreading out and forming a sheet. Analysis of the degree distribution of the cells in the tissue (Fig. 4c) shows a similar distribution to that recovered from the cell membranes in figure 2c. The tissue contains multiple cell-types necessary for maintaining the tissue homeostasis (Fig. 4d) [20]. A key question to understand the extrusion of cells from the zebrafish telencephalon is the spatial organization of cell phenotypes within the tissue. One method is to study the distribution and properties of clusters of cells of similar phenotype, i.e. a group of cells of the same cell type that are connected to each other. Such clusters can be identified by analyzing the network (Fig. 4e), which makes it possible to compute the proportion of cells of each type that belong to clusters of size larger than 1 (Fig. 4f). We can look at the distribution of cell clusters within the tissue to determine the statistical properties of the organization of the tissue (Fig. 4e).

Looking at the proportion of clusters strictly larger than 1, we see differences depending on the cell types. GFAP+/PCNA-cells, which compose 81% of the tissue, all belong to the same cluster. On the other hand, despite differing presence within the tissue, GFAP+/PCNA+ and GFAP-/PCNA+ cells have a similar chance of belonging to a cluster. Using the network representation makes it possible to investigate experimental data in a new light, whilst highlighting the cell-cell interactions within the samples.

## IV. DISCUSSION

Here we introduce `Griottes`, an open source, generalist algorithm that can generate the network representation of segmented images. The routine accepts a wide range of inputs: 2D and 3D images, mono-channel or multi-channel, as well as data tables. It is designed to be agnostic of the segmentation method used to pre-process the images.

By generating a graph representation of the input image, `Griottes` enables the use of a wide range of analysis tools, some of which are illustrated in the current paper. These include properties such as cell positions, geometry, fluorescence intensity levels, and contact surfaces with other cells in the tissue. The network representation then allows the geometric or phenotypic markers to be related among neighboring cells in order to understand the interactions that emerge.

The network representation of tissues condenses information contained in images. Instead of representing the system at the pixel-level, it summarizes the image at the single cell level, drastically reducing the memory uptake and increasing the information density. This analysis framework is particularly relevant when focusing on individual interactions between cells in the system and that these interactions are determined by the spatial properties of the tissue. Later developments will include information about the extra-cellular matrix or external constraints, which are simple to add to the network description.

`Griottes` is part of a growing ecosystem of imageanalysis tools enabling the spatial analysis of biological images [3, 17, 28, 29]. The library is built in Python and uses state of the art open-source libraries, allowing investigators to seamlessly integrate `Griottes` in their image and data analysis workflows. We expect to continue developing `Griottes`, with the long-term goal to make the step from pixel-based to node-based analysis as seamless as possible.

## CODE AND DATA AVAILABILITY

`Griottes` is an open-source Python package available in the following GitHub repository. Small files (e.g. CSVs) are accessible on the example repository, larger files (e.g. images) are available upon request. All the data are drawn from Dray et al. [20] and from unpublished lab experiments.

## METHODS

### MSC spheroid preparation

Human Mesenchymal stem cells (hMSCs) derived from the Wharton’s jelly of the umbilical cord [American Type Culture Collection (ATCC) PCS-500-010, LGC, Molsheim, France] were obtained at passage 2. The cells were maintained as previously described [24]. To isolate the CD146dim and CD146bright subpopulations, hMSCs (at passage 5) were recovered from the flasks using TrypLE. Then, the cells were incubated with a staining solution containing a 1:100 mouse anti-human CD146-Alexa Fluor 647 antibody (BD Biosciences) diluted in 1% FBS, for 30 min. 25% of the brightest and 25% of the dimmest CD146 stained cells were isolated by flow cytometry using a FACSAria III (BD Biosciences). The isolated CD146dim cells were then labelled with VybrantTMDiO, while the CD146bright cells were stained with VybrantTMDIL, for 30 min at 37 °C, as previously described [24]. After PBS washing, 6 × 106 of a mix of 50:50 CD146dim and CD146bright sorted-hMSCs/mL were seeded together into a 384 wells plate, which were each filled with 50 uL culture medium containing 1 uM SiR-DNA (Spirochrome), and the cells were let to form spheroids for 24 hours. The spheroids were then fixed using a 4% paraformaldehyde (PFA) solution (Alpha Aesar) for 30 min, and, after PBS washing, the aggregates were mounted into an agarose gel (Sigma) for 3D imaging.

### Imaging

Light sheet imaging has been performed using a commercial version of the Dual Inverted Single Plane Imaging Microscope (DiSPIM) [30] (Marianas −3i-). This system consists in two identical arms fitted with illumination and detection apparatus for light sheet generation and its subsequent imaging.

Despite its ability to acquire the same volume on two different angles with 90 degrees rotation, spheroids samples were acquired using only a single side acquisition. This provided acceptable signal to noise ratio for adequate segmentation and spared the time consuming dual view reconstruction.

The used objective was a water immersion 40x objective with 0.8 numerical aperture. This provides a resolution of 0,162 um/pixel on the fitted camera (Hamamatsu Orca Flash 4.0). Fast single plane acquisition was performed by objective translation via a piezo-electric element over a 150 um by 0.5 um increment. This provided a final cubic volume of 248 x 248 x 150 um of the acquired spheroid.

In order to excite the labelled samples, each plane was excited with 100 ms (Camera exposure time) sequential illumination of three discrete wavelengths: 488, 561 and 640 nm corresponding to GFP, RFP and Sir-DNA respectively. Chromatic fluorescence selection is done via a set of quad-band filters and dichroics (Semrock DA/FI/TR/Cy5-4X-B) on the CMOS Camera.

## ACKNOWLEDGMENTS

The authors thank Nicolas Dray and Laure Bally-Cuif for access to the zebra-fish images and useful discussions. We gratefully acknowledge the UtechS Photonic BioImaging (Imagopole), C2RT, Institut Pasteur, supported by the French National Research Agency (France BioImaging; ANR-10–INBS–04; Investments for the Future). We thank the support of the Inception program (Investissement d’Avenir grant ANR-16-CONV-0005). The DiSPIM microscope system benefited from financial support provided by the Region Ile de France via the DIM Elicit program. We also acknowledge the cytometry and biomarkers platform (CB-Utechs) for use of the cell sorters.

